# Lateralized foraging induces asymmetric corticostriatal plasticity in mice

**DOI:** 10.64898/2025.12.30.696996

**Authors:** Kenza Amroune, Maud Schaffhauser, Thomas Morvan, Ingrid Bureau, David Robbe

## Abstract

With practice, animals perform reward-oriented actions faster and with less variability. The dorsal striatum (DS) plays a key role in this process, potentially through opposing changes in cortical input strength to the two main striatal projection neurons (D1- and D2-SPNs). To test this hypothesis, we trained mice in a foraging task requiring them to perform quarter-turns (QTs) in a single direction (counterclockwise, CCW) along the walls of square towers to collect drops of water. As training progressed, the number and speed of CCW QTs increased while the variability of their trajectory decreased. Whisker trimming in well-trained mice altered QTs kinematics, highlighting the role of tactile inputs in guiding these actions. Combining ex vivo patch-clamp recordings in the DS with glutamate uncaging in the barrel cortex, we mapped the cortical neurons monosynaptically connected to D1- or D2-SPNs and measured the strength of these connections in the contralateral hemispheres, relative to the turn direction. In well-trained mice, compared to naive controls, the contralateral hemisphere showed no major changes in D1- or D2-SPNs connectivity, despite increased excitation of cortical pyramidal neurons. In contrast, the ipsiversive hemisphere exhibited no change in cortical excitability but showed increased cortical connectivity and input strength to SPNs. These findings reveal an unexpected hemispheric asymmetry in corticostriatal connectivity during lateralized foraging, which may reflect a homeostatic process normalizing striatal activity across hemispheres despite unbalanced cortical input.

## Introduction

Animals modify the way they execute actions leading to rewards as they associate specific sequences of movements performed in a given environment with the systematic obtention of a particular reward. This modification occurs both in time (actions are performed faster) and in space (their trajectories become less variable) (Jin and Costa, 2010; Dhawale et al., 2019; Lemke et al., 2019; Safaie et al., 2020; Dhawale et al., 2021; Mizes et al., 2023; Schaffhauser et al., 2025). Many studies have provided evidence that the dorsal striatum (DS) contributes to the kinematics of reward-oriented movements (Kim et al., 2014; Panigrahi et al., 2015; Rueda-Orozco and Robbe, 2015; Yttri and Dudman, 2016; Sales-Carbonell et al., 2018; Hidalgo-Balbuena et al., 2019; Jurado-Parras et al., 2020; Dhawale et al., 2021; Mizes et al., 2023; Park et al., 2025; Zheng et al., 2025). Additionally, striatal neuronal activity changes as animals progressively automatize reward-oriented behaviors (Barnes et al., 2005; Jin and Costa, 2010; Rueda-Orozco and Robbe, 2015; Sippy et al., 2015, 2021; Peters et al., 2021; Zareian et al., 2023) and coordinated neural activity emerges across the cortex and striatum during learning (Koralek et al., 2013; Lemke et al., 2019). Moreover, several studies suggest that the plasticity of excitatory cortico-striatal (CS) synapses is necessary for learning of reward-oriented behaviors (Jin and Costa, 2010; Koralek et al., 2012; Santos et al., 2015; Xiong et al., 2015; Lemke et al., 2021; Hwang et al., 2022). Still, whether such changes affect similarly the two main types of projection neurons in the DS remains unclear.

SPNs can be broadly divided into two main populations: D1-SPNs, which express dopamine D1 receptors, and D2-SPNs, which express dopamine D2 receptors (Kreitzer and Malenka, 2008) but see (Bonnavion et al., 2024). These neurons project to distinct targets within the basal ganglia, forming two functionally opposing pathways: the direct pathway (D1-SPNs), known to facilitate movement, and the indirect pathway (D2-SPNs), which suppresses it (Mink, 1996; Kravitz et al., 2010). The opponent modulation of movement following activation of D1- and D2-SPNs is well established in the context of locomotion and orienting-movements: unilateral prolonged activation of D1-SPNs induces contraversive turns (i.e., opposite to the activated hemisphere), while activation of D2-SPNs produces ipsiversive turns (Kravitz et al., 2010; Tecuapetla et al., 2014; Cregg et al., 2024). In contrast, during spontaneous locomotor activity, D1- and D2-SPNs are co-activated around action initiation, irrespective of action type (Tecuapetla et al., 2014; Barbera et al., 2016; Klaus et al., 2017; Markowitz et al., 2018; Parker et al., 2018). Still, several studies have revealed that, contrary to spontaneous exploratory activity, learned goal-directed behaviors depend on distinct, when not opponent, function of D1- and D2-SPNs (Hikida et al., 2010; Kravitz et al., 2012; Tai et al., 2012; Sippy et al., 2015; Yttri and Dudman, 2016; Nonomura et al., 2018; Cruz et al., 2022; Lowet et al., 2025) but see (Cui et al., 2013). This raises the possibility that well-learned lateralized movements toward rewards (e.g., always taking a left turn in a particular location to reach a goal) could be associated with different levels of activity of these two types of neurons, brought by asymmetric corticostriatal synaptic plasticity during reinforcement. This hypothesis has not been tested, primarily due to the lack of tasks requiring the performance of lateralized reward-oriented turns.

We recently characterized features of the functional connectivity between the barrel cortex and SPNs and found no major difference in the way this sensory region connects D1- and D2-SPNs in naive mice (Amroune et al., 2025). To probe whether learning of a lateralized reward-oriented sensory-guided action is associated with opponent changes in connectivity between sensory cortex and D1/D2-SPNs, we used a novel patch foraging paradigm (Schaffhauser et al., 2025) in which mice were required to make counterclockwise (i.e., lateralized) quarter-turns (CCW QTs) along the walls of squared towers to obtain drops of water. Based on the known ability of D1- and D2-SPNs to oppositely regulate turn direction, we hypothesised that, in the hemisphere contraversive to the rewarded turn direction, task learning would induce opposite changes in the connectivity and/or synaptic strength of inputs from the barrel cortex toward SPN, with an increase in inputs to D1-SPNs and/or a decrease in inputs to D2-SPNs (**Fig. 1**).

**Figure 1.**
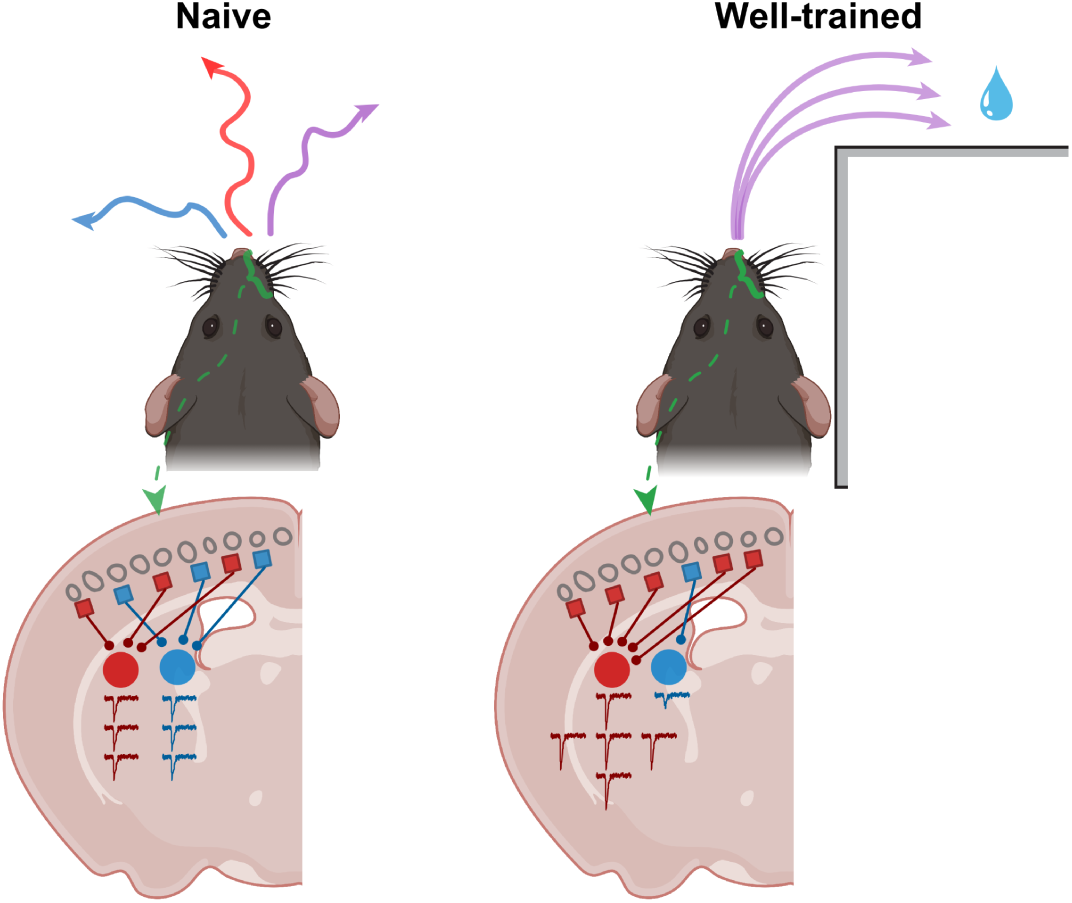
Schematic of the central hypothesis. **Left:** In naive animals, corticostriatal inputs are similar between the two striatal pathways: D1-SPNs (red) and D2-SPNs (blue). **Right:** With the learning of reward-oriented lateralized actions, such as rightward turns, corticostriatal plasticity would emerge, characterized by strengthened inputs to D1-SPNs (red) and weakened inputs to D2-SPNs (blue) in the contralateral hemisphere.

Within a few training sessions, mice performed the task proficiently, showing a selective increase in the number, speed and stereotypy of rewarded CCW QTs. The speed and stereotypy of CCW QTs were reduced following trimming of the whiskers, confirming their contribution to turn execution (but not turn selection). Using an ex-vivo slice preparation from naive or well-trained mice, we found no significant differences in the number, organization or strength of cortical inputs to D1- or D2-SPNs between trained and naive animals in the contralateral hemisphere (relative to turn direction and sensory stimulation). However, excitability of pyramidal neurons in the contralateral barrel cortex was increased in trained mice. The ipsiversive hemisphere, which was less stimulated during unilateral foraging, showed no increase in cortical excitability but an increase in cortical connectivity to SPNs. Altogether, our findings reveal an unexpected hemispheric asymmetry in corticostriatal plasticity which may reflect a homeostatic process normalizing striatal activity across hemispheres despite unbalanced cortical input.

## Results

### Mice quickly learned a whisker-guided laterized foraging task

We used a novel foraging task in which freely moving mice could obtain water rewards delivered by water ports protruding from the walls of 4 squared towers (Schaffhauser et al., 2025). Specifically, rewards were only delivered when water-restricted mice performed CCW QTs along the walls of the towers (e.g., from the east spout to north spout protruding from a given tower, **Fig. 2A**). Each time a mouse approached a given tower, a maximum number of available rewards was randomly selected between 4 and 12. When this number was reached, water drops were no longer delivered even if the mice performed additional CCW QTs (i.e., the tower was depleted). Mice were free to leave a tower before its full depletion (**Fig. 2A**). To encourage thigmotaxis, the task was conducted in complete darkness.

**Figure 2.**
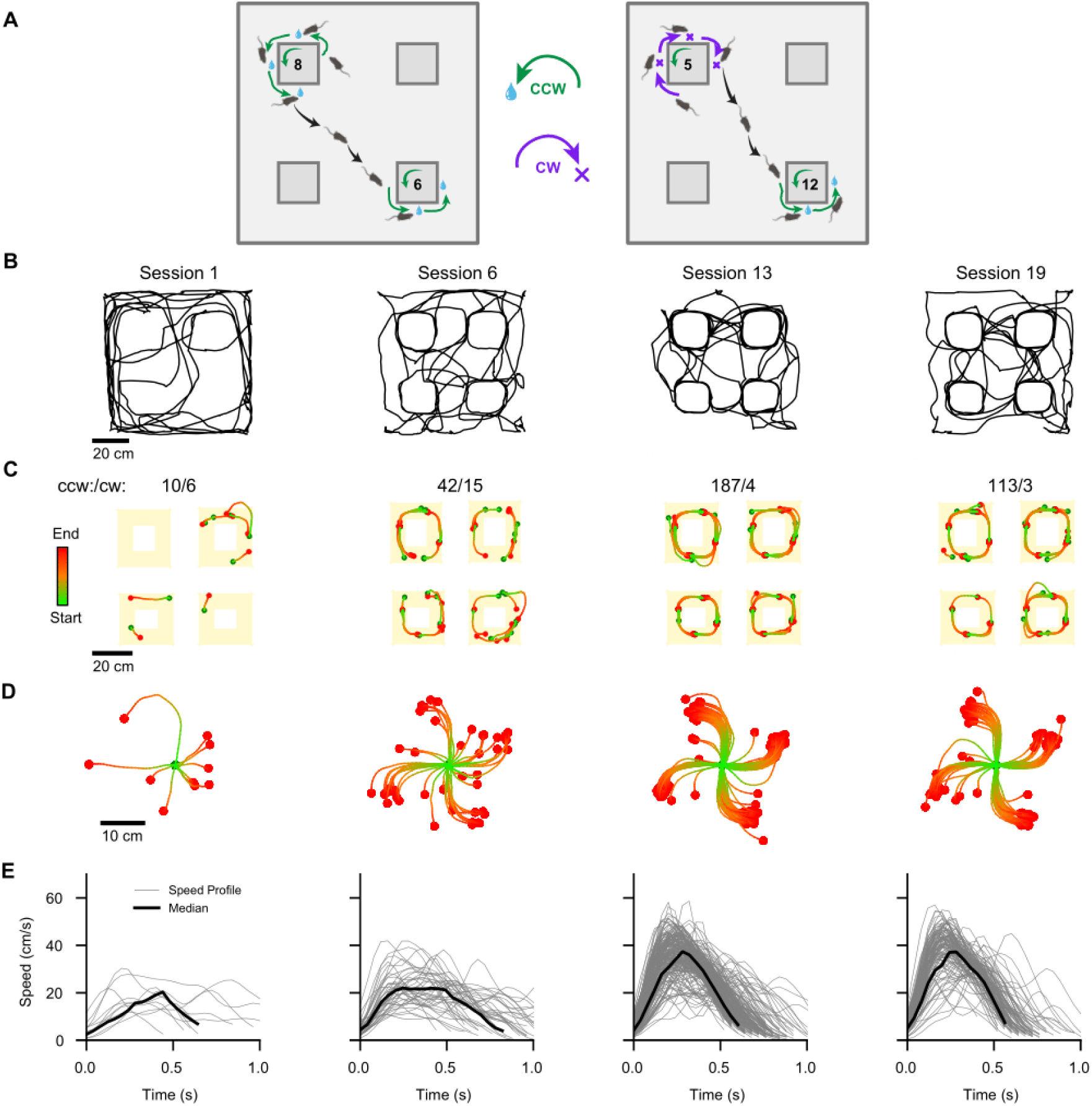
Increased efficiency of a single mouse performing a lateralized foraging task. **A.** Schematic of the task design. Counterclockwise (CCW) turns (green) around towers are rewarded with water, while clockwise (CW) turns (violet) are not rewarded. The number of maximum consecutive rewards for each tower is randomly picked between 4 and 12 consecutive quarter turns rewarded (8 and 6 in the first example, 5 and 12 in the second). **B.** Representative trajectories of a single animal during four different sessions (Sessions 1, 6, 13, and 19). Trajectories show movement paths during the first four minutes of each session. **C.** Up: Number of CCW and CW turns for each session. Bottom: Representation of CCW and CW turns around the 4 towers (yellow) for the same mouse and sessions as in (B). Turns trajectories are color-coded, with green indicating the start and red the end of the movement. **D.** All the CCW trajectories around towers from a single session are realigned relative to their origin. Turns trajectories are color-coded, with green indicating the start and red the end of the movement (same as in B and C). **E.** Speed profile of CCW turns for the same sessions as in (B), (C), and (D). Black lines represent the median speed of CCW turns.

Examining the trajectory of a single mouse revealed that during the first session, as expected, this animal primarily ran along the external walls of the arena (**Fig. 2B**, left). However, across sessions this animal spent progressively more time running around the towers (**Fig. 2B**, right). To quantify this change in behavior, we detected the turns made by the mouse to go from one water spout to the next one (quarter-turn around a tower corner, QT; Methods; **Fig. 2C**). Across sessions, more QTs were performed in the rewarded direction (CCW), but not in the unrewarded one (CW, **Fig. 2C**). Moreover, when aligning QTs to their starting points within each session (**Fig. 2D**), a decrease in trajectory variability was visible. Finally, examining the speed profiles of the QT in the rewarded direction revealed a prominent increase across sessions (**Fig. 2E**). Altogether the increase in number of CCW QT and changes in speed and variability are congruent with this mouse maximizing its harvesting rate across sessions (Schaffhauser et al., 2025).

We next examined whether the behavioral changes observed in a single mouse were robust at the group level (**Fig. 3**, N=17, 6 females). Task performance was quantified across all sessions performed (range of sessions performed: 15-19, only the first 15 sessions are presented). We observed the same behavioral changes at the population level as described above, namely an increase in proportion and number of reward-oriented QT (CCW, **Fig. 3AB**), decreased trajectory variability of reward-oriented runs (**Fig. 3CD**), and increased speed (**Fig. 3DC**). Critically these kinematics effects were selective of reward-oriented turns (i.e., occasionally performed CW turns were slow and their trajectory variable). Finally, to quantify the extent to which mice exploited a tower before switching to another one, we computed the mean number of QTs performed per rewarded visit per session. For all the mice, this number increased across sessions but remained below the theoretical average of available rewards per visit (i.e., 8; **Fig. S1**), indicating that mice under-exploit the tower in this task. Altogether, these results show that mice subjected to this foraging protocol quickly developed lateralized and efficient reward-oriented full-body movements and a strategy (underharvesting) allowing them to efficiently accumulate drops of water.

**Figure 3.**
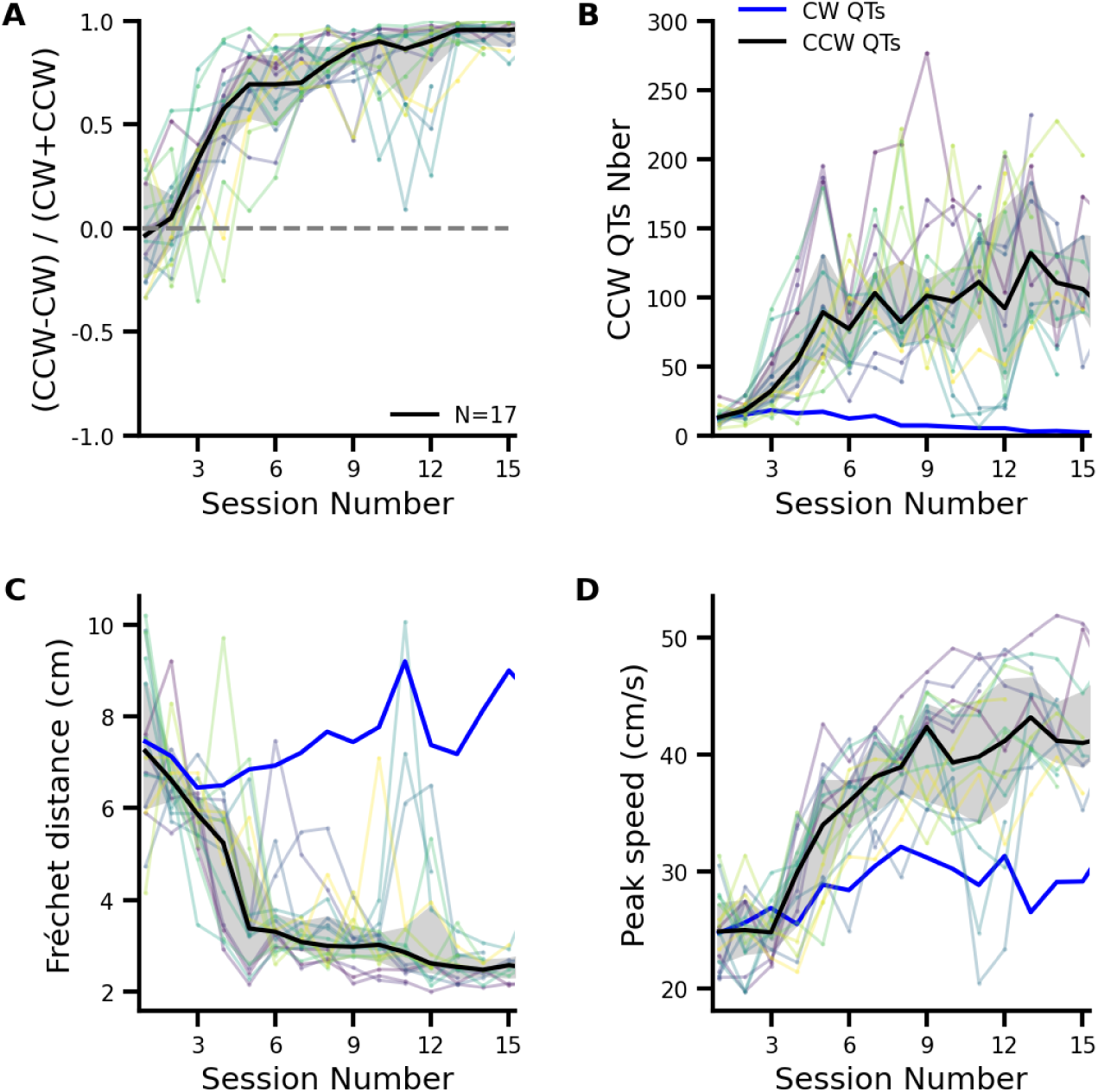
Mice quickly learn a lateralized foraging task and modified the kinematics of their reward-oriented actions. **A.** Normalized directional preference of QTs across sessions and all mice (N=17). Each colored line represents a single mouse, while the black line indicates the median across animals. The grey shaded area denotes the interquartile range (25th to 75th percentile). **B.** Number of CCW QTs per session. Each colored line represents a single mouse, and the solid black line indicates the median. The median number of CW QTs across sessions is represented in blue. **C.** Median Fréchet distance between pairs of CCW (black) and CW (blue) QTs trajectories (rotated and aligned relative to a single tower corner, see Methods) performed during each session. Lower values indicate less variable trajectories. **D**. The session-median maximum speed (cm/s) of CCW turns is shown for each mouse across sessions (black line indicates the group median). The median speed of CW turns across sessions is shown in blue.

As we hypothesized that the functional connectivity between the barrel cortex and the striatum changes as mice learn to perform efficient, reward-oriented turns, we first verified that they relied on their whiskers to execute these turns. For this, we trimmed their whiskers (N=6, methods) after 12 or 14 foraging sessions. When comparing the kinematics of CCW QTs in the session preceding and following whisker trimming (methods), reward-oriented turns became slower (**Fig. 4A**) and more variable (**Fig. 4B**). These alterations were not paralleled by a decrease in the number of CCW QTs (**Fig. 4C)** nor an increase in the number of QTs in the unrewarded direction (clockwise, CW) (**Fig. 4D)**, showing that the whisker trimming did not perturbate procedural learning or motivation but selectively contributed to the kinematics of reward-oriented actions. Finally, the brief exposure to isoflurane necessary to trim the whisker did not, by itself, induce comparable effects (**Fig. S2**). These results indicate that mice use whisker-dependent sensory feedback to constrain the kinematics of their automatized reward-oriented turns along the walls of towers.

**Figure 4.**
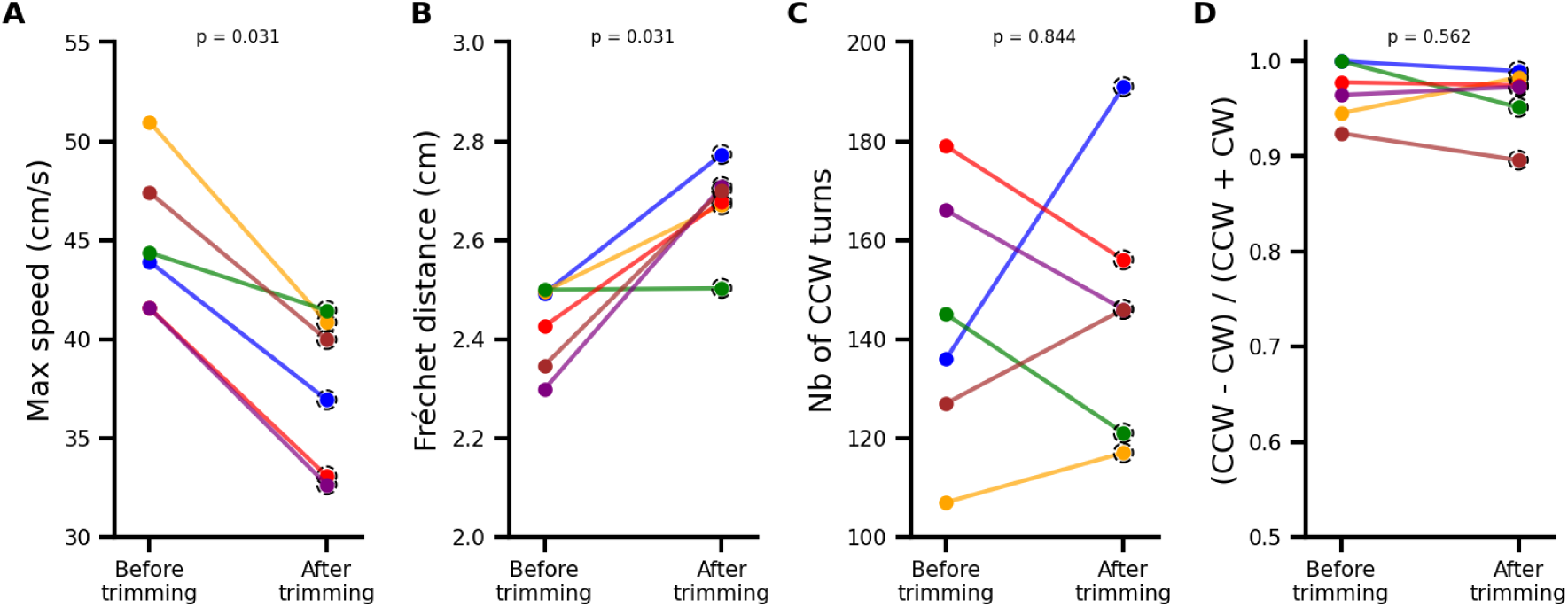
Foraging kinematics alteration following whisker trimming. **A.** Median max speed of CCW turns for individual mice before and after whisker trimming. Each colored line represents an individual. **B.** Fréchet distance (cm) of CCW turns for individual mice before and after whisker trimming. Lines represent individual mice, with colors matching (A) and (B). **C.** Number of CCW turns for individual mice before and after whisker trimming. Each colored line represents an individual. **D.** Ratio of CCW turns for individual mice before and after whisker trimming. Each colored line represents an individual. *p* values at the top of each panel are derived from Wilcoxon signed-rank test.

### CS Inputs to D1 and D2 SPNs Following Lateralized Learning

To explore if learning a lateralized reward-oriented action was associated with changes in CS projections to D1 and D2 SPNs (see predictions in **Fig. 1**), we mapped functional monosynaptic connections using a newly developed parasagittal ex-vivo preparation preserving the connectivity between several whisker-columns of the barrel cortex and SPNs neurons in the dorsolateral striatum (DLS, Amroune et al., 2025). We combined patch-clamp recordings of SPNs in whole-cell voltage-clamp configuration with glutamate uncaging through laser scanning photostimulation (LSPS), a stimulating method allowing us to determine the location of cells presynaptic to individual SPNs and the strength of their synaptic connections (**Fig. 5A**, left). Glutamate was uncaged over a 29 x 16 pixel grid (2.1 × 1.1 mm, 75 μm spacing) “placed” over the barrel cortex. Each time glutamate uncaging evoked an EPSC in the SPN, the location of the site and the amplitude of the EPSC were reported on a map, displaying the spatial origin and strength of its cortical projections (**Fig. 5A**, right). Animals were Drd1a-tdTomato hemizygous mice in which D1 receptor expressing neurons were labeled. Unlabeled neurons are principally SPNs expressing the D2 receptor in these mice (Bertran-Gonzalez et al., 2008; Ade et al., 2011; Enoksson et al., 2012; Thibault et al., 2013; Cao et al., 2018). We searched for changes in CS projections to D1- and D2-SPNs by examining three parameters contributing to cortico-striatal functional connectivity: the width of input field in the barrel cortex (**Fig. 5B**, left), the number of cortical columns connected to a single SPN (**Fig. 5C**, left), and the amplitude of EPSCs (**Fig. 5DE**, left). Despite the fact that all mice displayed a strong bias for CCW QT and executed them quickly and with little variability, no significant changes in the level nor in the horizontal organization of corticostriatal connections to either D1- or D2-SPNs were observed between naive and trained animals (**Fig. 5B-E**, center and right). Further analysis of EPSCs summed in each layer (2/3, 4 5a, 5b, 6) also showed no significant change for D1-or D2-SPNs (**Fig. 5FG**). Similarly, the proportion of cells receiving input from each cortical layer remained similar between naive and trained animals (**Fig. 5HI**), with a major contribution of upper layer 5 (∼50%) and a smaller one from the superficial layer (∼20% from layer 2/3), as previously described (Wise and Jones, 1977; Reiner et al., 2003; Wall et al., 2013; Yamashita et al., 2018; Amroune et al., 2025). A significant increase in the proportion of D2-SPNs receiving input from the layer 5a was nevertheless detected (**Fig. 5I**), without alteration of the strength of this projection compared to naive mice **(Fig. 5G)**. These results indicate that, in the contralateral hemisphere relative to the learned turn direction, the overall organization and synaptic strength of corticostriatal projections from the barrel cortex to both types of SPNs remain largely preserved following learning of a lateralized foraging task.

**Figure 5.**
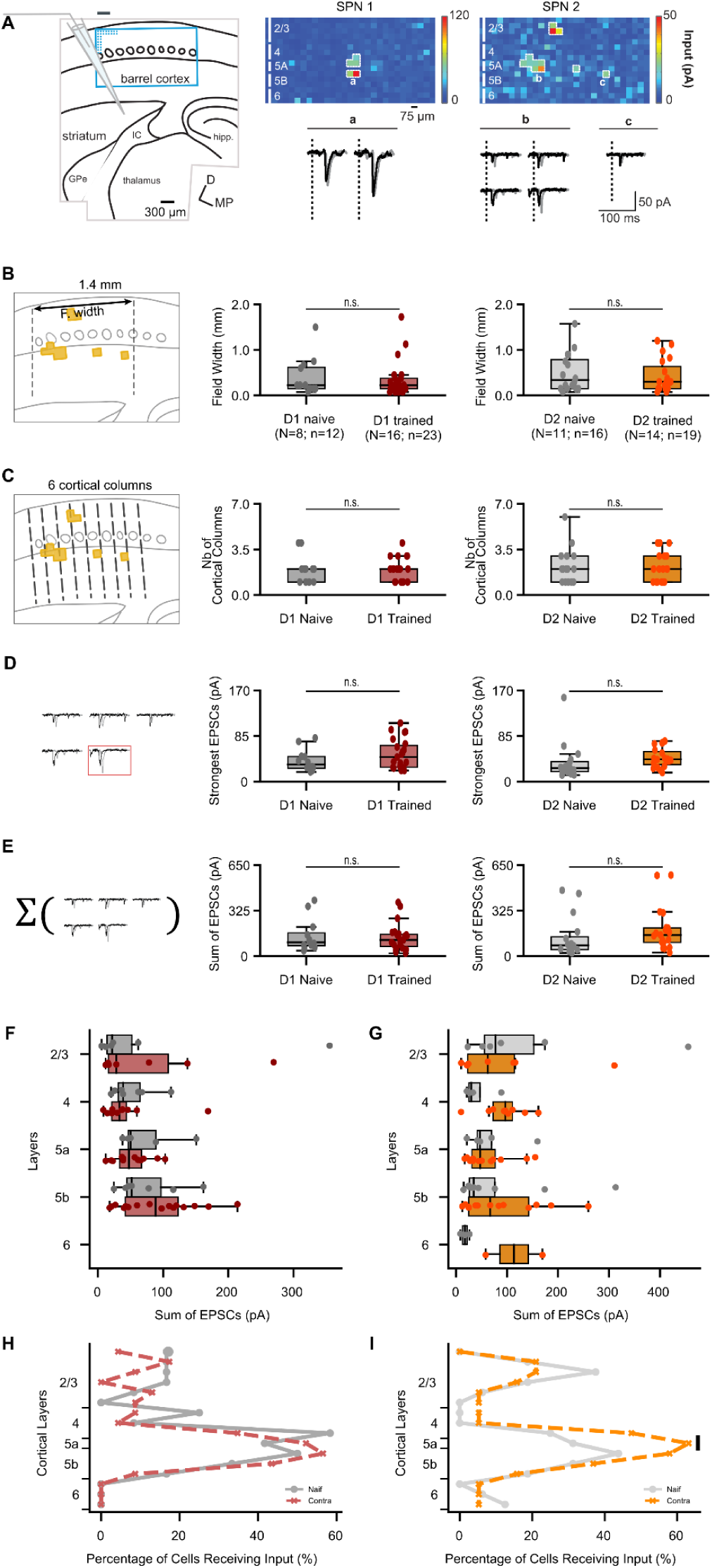
No differences between naïve and trained animals in the number and strength of cortical inputs to D1/D2 SPNs. **A.** Left: Layout of the experiment. A single SPN was recorded in the dorsolateral striatum while cortical neurons were photostimulated with LSPS. The grid of LSPS (blue) was positioned on the barrel cortex. GPe, globus pallidus, external segment; IC, internal capsule; hipp., hippocampus. Right Top : Synaptic input maps for two individual SPNs, labeled SPN 1 and SPN 2. SPN 1 shows a single cluster of input, whereas SPN 2 exhibits four distinct clusters. Each color pixel represents the peak amplitude of excitatory postsynaptic currents (EPSCs) within a 50 ms window following stimulus onset. Cortical layers are marked with solid white lines on the left side of each map, and inputs are highlighted with white dashed borders around relevant pixels. Right bottom: Example EPSCs recorded from the two SPNs. Responses were evoked by the photorelease of glutamate at the sites indicated by letters a-c in the maps shown above. Two repetitions are superposed (black and gray). Stimulus onsets are indicated by the vertical dashed lines (2 ms duration). **B-E.** No significant changes in the CS innervation of D1 or D2 SPNs after learning (D1, n=12 cells in N=8 mice; D2, n=16, N=11), in field width (mm, B); number of cortical columns (C); strongest excitatory postsynaptic currents (EPSCs, D) and sum of EPSCs (E). **F-I.** Laminar input distribution. The sum of EPSCs to individual SPNs as a function of their laminar origin (F-G). Panel H-I shows the contribution of each cortical layer to the SPN innervations. The 16 rows of the grid (Fig 5A) correspond to different cortical layers (Layer 2/3: 1-6; L4: 7-9; L5a: 10; L5b: 11-13; L6: 14-16). The vertical black line indicates a significant difference between the two groups (permutation test).

### Increased Cortical Excitation and Interhemispheric Asymmetry in CS Connectivity

One possibility is that modifications occurred upstream of the DLS. We examined whether barrel cortex layer 5 (L5) pyramidal neurons displayed change in their excitation, as they were the primary source of CS projections in our experiments (**Fig. 5HI**). To characterize this excitation, we recorded action potentials (APs) in current-clamp mode evoked by the uncaging of glutamate (**Fig. 6AB**). Interestingly, trained animals displayed a significantly higher number of APs in L5 pyramidal neurons compared to naive animals (**Fig. 6C**).

**Figure 6.**
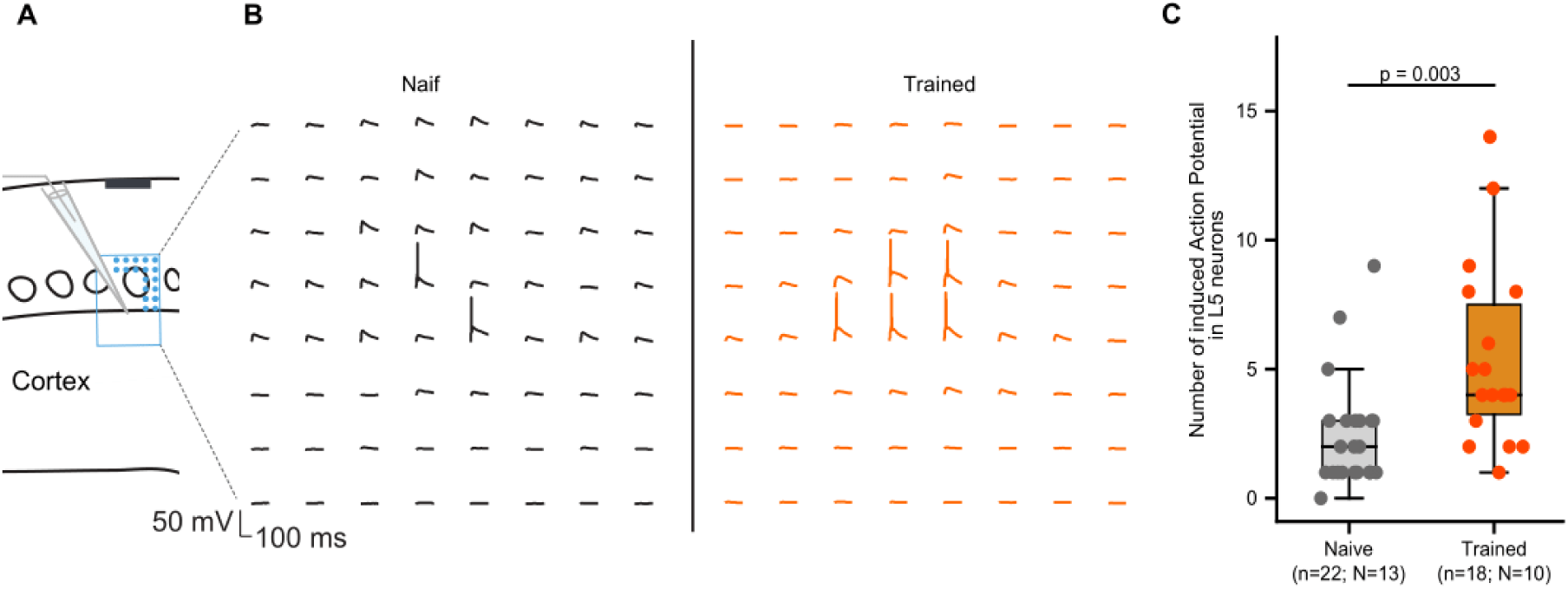
Increased excitation of cortical layer 5 neurons in the contralateral hemisphere, after training. **A.** Experimental layout showing cortical Layer 5 (L5) neuron recordings in the barrel cortex, while glutamate is uncaged around its soma and dendrites. **B.** Representative traces of action potentials (APs) evoked by glutamate uncaging in L5 neurons of the barrel cortex, recorded in naive and trained animals.The scale bar represents 100 ms and 50 mV. **C.** The number of evoked action potentials by glutamate uncaging in L5 neurons for naive (n=21) and trained (n=18) conditions. Each dot represents a cell. Trained animals show a significantly higher number of evoked APs compared to naive animals (p = 0.003).

To determine whether the increase in cortical excitation arose from enhanced contralateral whisker stimulation associated with lateralized foraging, or instead reflected a more global change in motivational state (i.e, which should affect both hemisphere), we assessed whether a similar increase was present in the less-stimulated ipsiversive hemisphere (**Fig. 7**). To preserve barrel cortex–DLS connectivity, our ex vivo preparation requires a mediolateral slicing angle, which prevents examining CS connectivity in both hemispheres of a single animal. We therefore trained a separate cohort of mice on the same foraging protocol, except that CW QTs were rewarded (**Fig. 7A**). In these animals, the ex-vivo preparation remained identical, but it was now taken from the hemisphere ipsilateral to the rewarded turn direction. L5 pyramidal neurons in the ipsiversive hemisphere exhibited significantly lower excitation than those in the contraversive hemisphere (**Fig. 7B–C**, p = 0.037), suggesting that the increased contralateral excitation in the barrel cortex resulted from the lateralized foraging. We then assessed whether the reduced excitation in the ipsilateral barrel cortex was associated with altered CS functional connectivity (**Fig. 7D**). Unexpectedly, SPNs in the ipsiversive hemisphere had wider input fields in the barrel cortex (**Fig. 7E**, p = 0.001), with a greater number of cortical columns contributing to these inputs (**Fig. 7F**, p = 0.004). This increased connectivity was particularly prominent in both superficial and deep layers, as SPNs in the ipsiversive hemisphere were more often innervated by upper L5 (86% vs. 57%) and superficial L2/3 (39% vs. 19%; **Fig. 7G**). We examined whether these changes in the number and convergence of cortical inputs were accompanied by alterations in the strength of individual connections. The amplitude of individual EPSCs remained similar between hemispheres (**Fig. 7H–I**). Thus, the observed increase in total EPSC sum in the ipsiversive hemisphere (**Fig. 7J**, p = 0.015) reflected a greater number of CS connections rather than changes in individual synaptic strength. Additionally, we noted greater variability in the total EPSC sum in the ipsiversive hemisphere, indicating more heterogeneity in input strength (**Fig. 7K**, p = 0.001). These unexpected results suggest that the ipsiversive hemisphere, while not over-stimulated during lateralized foraging, undergoes important changes leading to an overall increase in CS connectivity to individual SPNs.

**Figure 7.**
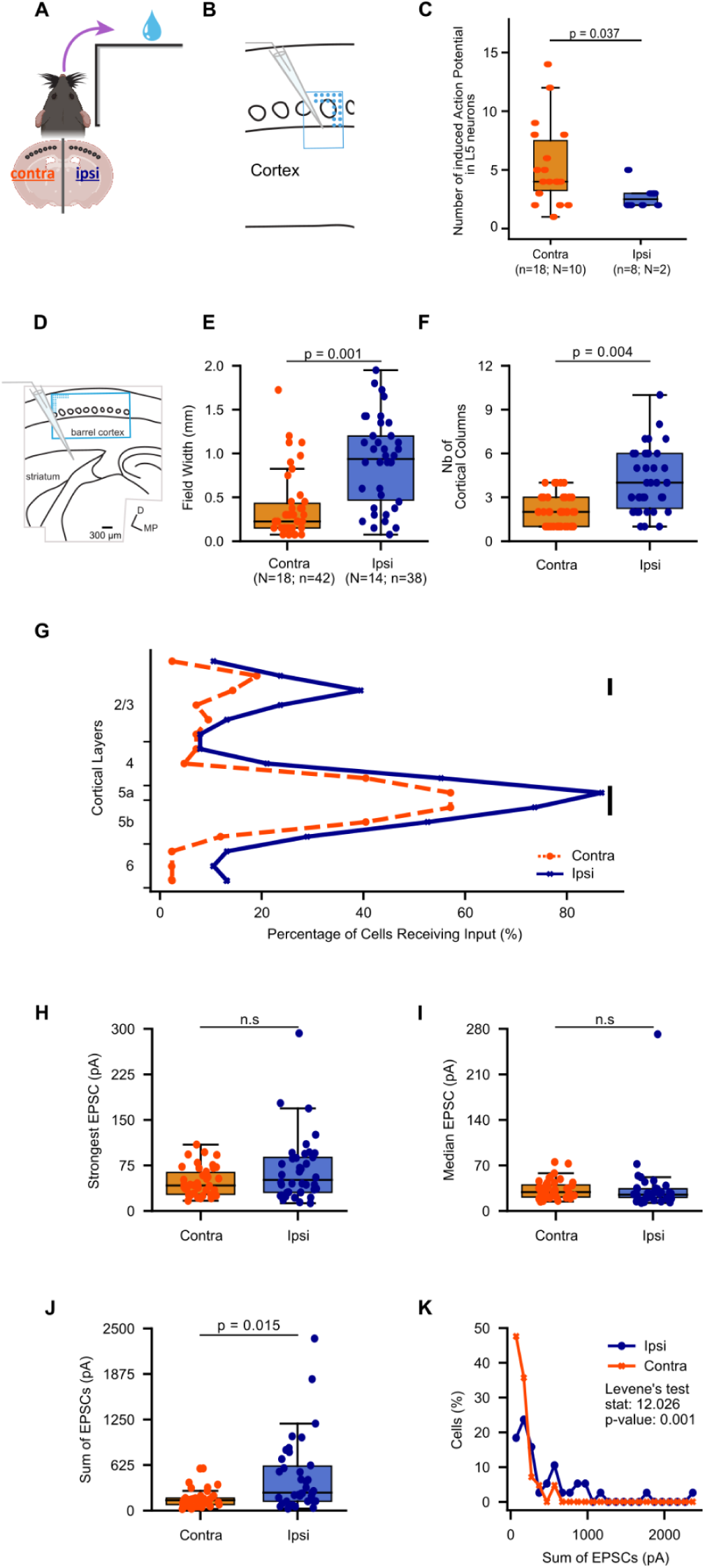
Increased functional connectivity between the barrel cortex and SPN in the ipsilateral hemisphere after lateralized foraging. **A.** Schematic of the task design. Animals were required to make CW QTs to study CS projections to the ipsiversive hemisphere. **B.** Experimental layout showing cortical Layer 5 (L5) neuron recordings in the barrel cortex, while glutamate is uncaged around its soma and dendrites. **C.** The number of action potentials (APs) evoked by glutamate uncaging in L5 neurons in the LSPS grid, for trained contraversive (n=18) and ipsiversive (n=8) conditions. Each dot represents a cell. The trained contraversive animals show a significantly higher number of evoked APs compared to ipsiversive animals (p = 0.037). **D.** Layout of the experiment for panels E-K. A single SPN was recorded in the contra or ipsi dorsolateral striatum while cortical neurons were photostimulated with LSPS. The grid of LSPS (blue) was positioned on the barrel cortex. **E.** Comparison of the field width formed by all the cortical neurons connecting single SPNs (see drawing Fig 5A) recorded in contralateral (Contra, n=42, N=18) and ipsilateral (Ipsi, n=38, N=14) hemispheres in trained mice. **F.** The same as in D for the number of columns connecting single SPNs. **G.** Contribution of each cortical layer to the SPN innervation. Vertical black lines indicate significant differences between the two groups (permutation test). **H-I.** Corticostriatal EPSCs show no significant differences between hemispheres, neither in the amplitude of largest responses (G, p = 0.2991) nor in the median of EPSCs (H, p = 0.2001). **J-K.** The sum of EPSCs and distribution show significant differences between hemispheres. The sum of EPSCs (I, p = 0.015). The distribution of sums of EPSCs, representing the difference in the variance between the 2 groups (I; p = 0.001, Levene test).

## Discussion

In this study, we used a novel foraging task to test whether learning a lateralized sensory-guided reward-oriented action (quarter-turns along the walls of towers to collect rewards) was associated with an increased (decreased) strength in functional connectivity between the barrel cortex and D1-SPNs in the contralateral hemisphere (D2-SPNs, **Fig. 1**). All the mice that were subjected to our foraging protocol developed fast and stereotyped reward-oriented movements with a strong directional bias, allowing them to efficiently obtain drops of water. Critically, the kinematics changes were altered following whisker trimming showing that mice used their whisker to optimize the kinematics of their movements (but not to choose which direction to turn). *Ex-vivo*, we investigated the excitatory projections from the barrel cortex to D1- and D2-SPNs in the hemisphere contralateral to the rewarded turn direction, by combining laser-induced glutamate uncaging and patch-clamp recording, an approach giving access to both the localization of presynaptic cortical neurons and the strength of their synapses on the SPNs. When comparing the CS projections between brain slices obtained from animals either trained at least in 15 foraging sessions or untrained, no major differences were observed in the hemisphere predominantly “stimulated” when mice performed reward-oriented actions (i.e., the hemisphere contralateral to the tower walls). Still, we observed increased excitation of the striatal-projecting layer 5 neurons in the barrel cortex. No such increase was observed in the ipsilateral barrel cortex (i.e., the hemisphere less “stimulated”). Surprisingly, this lower excitation of cortical neurons in the less-stimulated brain hemisphere was accompanied by an increase in connectivity between the barrel cortex and SPNs. Thus, a form of hemispheric adaptation occurred in mice performing lateralized sensory-guided turns. This suggests an interhemispheric homeostatic process that balances activity levels between the left and right striatum, independent of SPN subtypes.

Building on the well-known opposite modulations of orienting movements following unilateral perturbation of D1-SPNs and D2-SPNs (Kravitz et al., 2010; Tecuapetla et al., 2014; Bay Kønig et al., 2019), and on the other hand, the many studies reporting that D1 and D2-SPNs have opposite functions in the control of reward-oriented learnt behavior (Hikida et al., 2010; Kravitz et al., 2012; Tai et al., 2012; Sippy et al., 2015; Yttri and Dudman, 2016; Nonomura et al., 2018; Cruz et al., 2022; Lowet et al., 2025) but see (Cui et al., 2013) we tested the straightforward hypothesis that mice well-trained to perform sensory-guided turns in a single direction would lead to increased (decreased) corticostriatal connectivity on D1-SPNs (D2-SPNs), compared to naïve mice. However, no such changes were observed, as naive and trained mice displayed similar levels of cortical input in both cell types, visible in the number of connecting cortical columns and in the strength of projections. When considering the reasons for the absence of opponent changes in CS strength on D1- and D2-SPN, a first potential issue is whether our training protocol was strong enough to induce plasticity. Our data argue against this concern, as training not only induced detectable changes in the excitation of cortical neurons projecting to the DLS in the contralateral hemisphere (relative to whisker stimulation) but also increased CS connectivity in the less-stimulated hemisphere. A potential limitation of our study is that in the present training protocol, although mice did massively prefer CCW over CW turns, they under-exploitated the towers, leaving them before they were depleted (Figure S1 and Schaffhauser et al., 2025). D1-SPN and D2-SPN have been implicated in controlling the exploration/exploitation tradeoff, the latter favoring explorative behaviors (Humphries et al., 2012; Dunovan and Verstynen, 2016). In this regard, it is interesting that the only significant difference of CS connectivity on D1-SPN vs D2-SPN when comparing naive and trained animals is a small but significant increase in the proportion of D2-SPNs connected by layer 5a neurons. Future studies should therefore investigate whether manipulating bidirectionally the exploration-exploitation tradeoff (which was beyond the scope of this study) is associated with an opponent effect on the functional CS connectivity D1-SPNs and D2-SPNs.

Several previous studies have successfully used *ex-vivo* LSPS input mapping to detect changes in functional connectivity following sensory deprivation or conditioning (Shepherd et al., 2003; Rosselet et al., 2011; Jacob et al., 2012). This is the first time that such an investigation is made to detect plasticity of cortical inputs onto SPNs and one potential concern is that, on the contralateral side to whisker stimulation, the plasticity processes underlying the kinematic changes of specific actions might be confined to a subset of neurons, which would require more targeted recordings to detect. Indeed, although this has not been directly examined in our task, studies of self-paced locomotion in open fields have shown that nearby SPNs can be selectively recruited during specific actions (Barbera et al., 2016; Klaus et al., 2017). Because our SPN recordings were blind to their task-related activation levels, we cannot rule out the possibility that opponent CS synaptic changes occurred in subpopulations of D1 or D2 SPNs that we did not systematically sample. This caveat is nevertheless mitigated by a previous study showing robust pathway-specific plasticity in the DLS after a week of training on the accelerating rotarod (Yin et al., 2009).

Although we did not observe opponent changes in CS projections to D1- and D2-SPNs in the contralateral DLS of mice subjected to our lateralized foraging task, we did find an increased excitation in L5 pyramidal neurons, the primary source of cortical inputs to the striatum (Wise and Jones, 1977; Reiner et al., 2003; Shepherd, 2013; Wall et al., 2013). Specifically, more spikes were evoked by glutamate uncaging in cortical neurons taken from trained animals, compared to naive animals. This heightened excitation aligns with previous findings indicating that sensorimotor learning enhances cortical activity in sensory regions involved in task-related processing (O’Connor et al., 2010; Sachidhanandam et al., 2013; Yamashita and Petersen, 2016). Following associative sensorimotor learning, *in vivo* recordings in the DLS have highlighted an increase in sensory-evoked responses in SPNs (Sippy et al., 2021) and in sensory CS synapses (Xiong et al., 2015), leading us to hypothesize that whisker-related CS inputs to SPNs are strengthened in mice trained in the foraging task. While we did not find direct changes in the strength or number of CS connections, the increased excitation in L5 suggests an enhanced cortical activity that could still lead to more effective sensory activation of SPNs in the DLS, although whether this increased excitation is selective for neurons targeting one SPN subtype remains speculative in our study (Klug et al., 2025). Therefore, functional changes at the cortical level may be driving the behavioral adaptations and striatal activity changes observed after motor learning (Xiong et al., 2015; Sippy et al., 2021). This increase in pyramidal neuron excitation was specific to the hemisphere that was primarily stimulated during our lateralized foraging task (i.e., cortical excitation in the ipsilateral hemisphere was weaker compared to the contralateral one; **Fig. 7C**). Unexpectedly, we observed an increase in CS functional coupling in slices taken from this less-stimulated hemisphere. This heightened CS connectivity in the ipsilateral DLS may be a form of compensatory plasticity or a homeostatic adaptation, to normalize CS activity between hemispheres. The concept of homeostatic plasticity suggests that neural circuits can maintain stable activity by adjusting synaptic strengths to counterbalance changes in network activity (Turrigiano, 2008; Wen and Turrigiano, 2024). Therefore, increasing CS inputs to SPNs in the ipsiversive striatum might help stabilize SPN activity across hemispheres. One possible explanation for our findings is that the contraversive hemisphere, actively engaged in task performance, relies on enhanced cortical excitability to modulate the kinematics of reward-oriented movements, while the ipsiversive hemisphere compensates by increasing its synaptic input to maintain overall neural balance during other exploratory movements. Thus, highly lateralized motor learning involves not only neuronal changes within the contralateral hemisphere involved with processing sensory stimuli and modulating movements, but also compensatory adjustments in the ipsilateral one, indicating a more nuanced function of corticostriatal plasticity than previously thought.

In summary, our study failed to reveal opponent changes in CS connectivity on D1- and D2-SPNs during lateralized motor learning, but reveals unexpected asymmetric corticostriatal plasticity across hemispheres. We propose that these findings reflect the importance of homeostatic adjustments to maintain balanced excitation levels across the left and right striatal activity.

**Figure S1.**
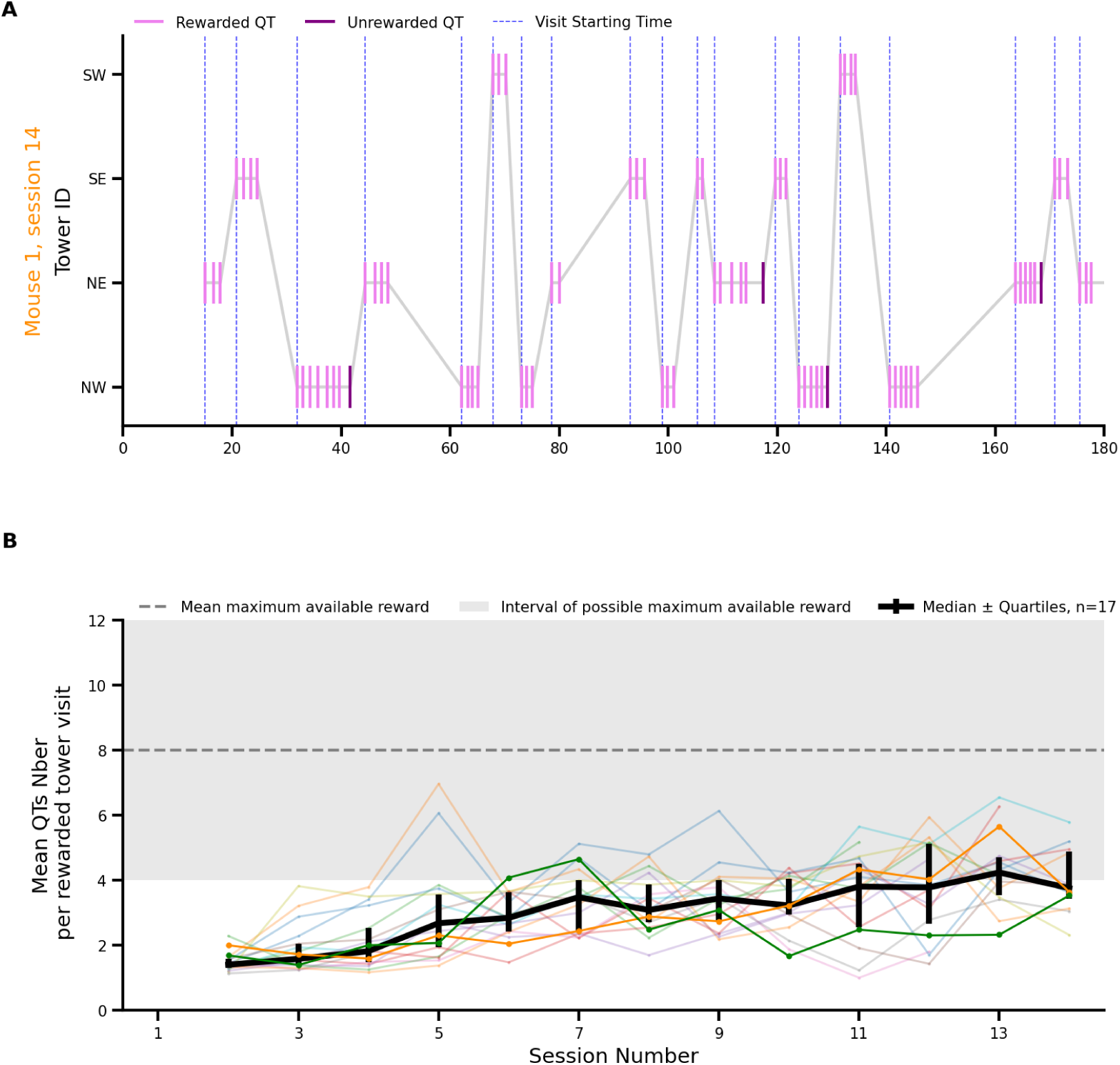
Mice performed short exploitation bouts. **A.** Sequence of QTs sorted by tower in which they occur for the same example mouse as in Fig. 2. **B.** Mean number of QTs per rewarded visit (i.e., visits with at least one rewarded QT) across sessions for all mice.

**Figure S2.**
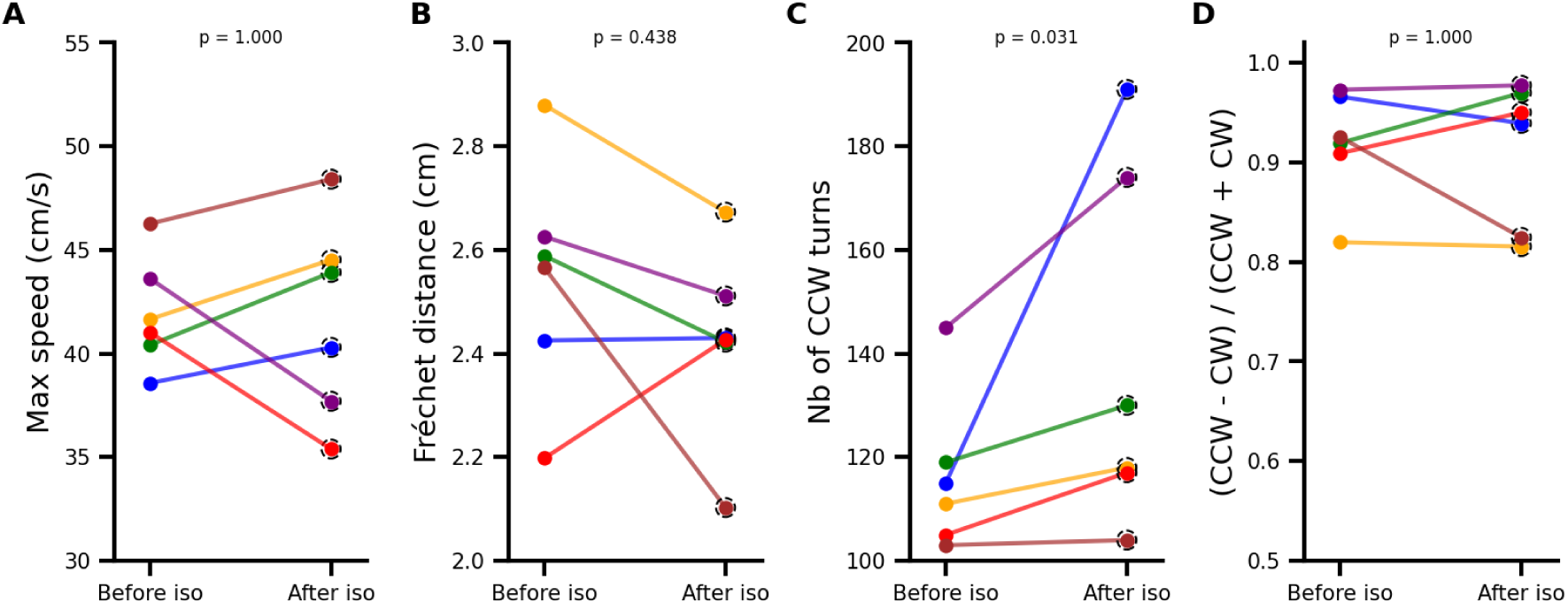
Brief isoflurane anesthesia does not impair quarter-turns directions and kinematics. **A.** Median max speed of CCW quarter-turns for individual mice before and after brief isoflurane anesthesia. Each colored line represents a single mouse. **B-D** Similar to A but for Fréchet distance of CCW quarter-turns (**B**), number of CCW QT quarter-turns (**C**), and normalized preference for CCW quarter-turns (**D**). *p* values at the top of each panel are derived from Wilcoxon signed-rank test.

## Methods

### Animals

A total of 37 hemizygous mice B6.Cg-Tg(Drd1a-tdTomato)6Calak/J mice (JAX stock #016204; (Ade et al., 2011) were used, both males and females (postnatal day [PND] 30–45). They were housed in groups of 2–3 mice on ventilated racks controlled by temperature (21 °C). A 12 h–12 h light/dark cycle was maintained and all experiments were carried out during the dark phase. Food was available *ad libitum* in their home cage, but mice had to perform the behavioral task described below to obtain water. Body weight was measured twice a day, before and after behavioral experiments. All experiments were performed under the European Union Directive (European Community Directive 86/609/EEC) and approved by the national ethics committee (Ministère de l’enseignement supérieur et de la recherche, France, Authorization # 46285).

### Task Apparatus

The experimental apparatus consists of a square arena (83 cm per side) enclosed by 9 cm high walls, with four smaller square towers (12 cm per side, 9 cm in height) positioned at the center of each quadrant (Figure 1) (Schaffhauser et al., 2025). Each of the 4 walls (north, east, south, and west) of the 4 towers (northwest, northeast, southwest, and southeast) is equipped with a water spout located at its center, 2 cm above the floor. Each spout extends 2 cm outward from its wall and is connected to a water reservoir housed within the tower, inaccessible to the mice (an opaque roof covers the arena and the towers). Single water drops (2–5 µL) can be delivered at each spout by transiently opening a solenoid valve (VAC-20 PSIG, Parker) placed between the reservoir and the spout (i.e. the apparatus contains 16 independently controllable solenoid valves).

The arena floor is made of transparent plexiglass, elevated 140 cm above the ground, allowing video tracking of the mice’s movements using an infrared camera placed beneath the apparatus (Grasshopper3 GS3_U3_41C6NIR, Point Grey Research, 25 Hz). The experiments are conducted in complete darkness. Video frames are processed online using a background subtraction algorithm provided by the open-source OpenCV library, integrated into a Python function. This function handles data acquisition and controls the solenoid valves based on the animals current and past positions. The Python function communicates with a Raspberry Pi, which independently controls each solenoid valve and triggers their transient opening when mice cross boundaries between predefined regions of interest (north, east, south, and west trapezoids surrounding each tower). The code generates four output files: 1) a raw video file; 2) a parameter file containing all experimental metadata; 3) a position file with the centroid position and time for each detected frame; 4) a “turn info” file containing detailed information about detected transitions between contiguous trapezoids, including their time, tower ID, starting trapezoid ID, landing trapezoid ID, whether the transition was rewarded.

### Behavioral task

In this version of the task, mice can obtain rewards by turning around any of the 4 towers from one spout to the next one (i.e., by executing quarter-turns, QTs) with the two following constraints: 1) only QTs performed in a counterclockwise (CCW) from the viewpoint of the camera were rewarded. This rewarded direction was fixed across all the sessions; 2) each time a mouse starts harvesting a tower, a maximum number of consecutive rewarded QTs is randomly drawn (between 4 and 12, corresponding to one and three full turns of tower) after which the current tower stops delivering reward and the mice must leave to exploit one of the three other towers. When a mouse leaves a tower, its consecutive number of rewarded turns is reset to zero and the tower will become active the next time the mice start harvesting it. The full training protocol is as follows: prior to training, mice were handled 2-3 times per day for 30 minutes, for 3 days. Following handling, the mice were familiarized with the arena through 1 session of 12 minutes (free exploration). After the familiarization session, the mice were weighed and placed on water restriction. The next session, training began and each CCW QT was rewarded. This initial learning phase (Phase 1, free exploitation) was carried out over 2 days (2∼3 sessions of 12 minutes per day separated by at least 3 hours). To encourage the usage of their whiskers, following the initial 2 days of testing in Phase 1, the number of consecutive rewards available around a tower was randomly set between 4 and 12 each time the mice started harvesting a tower. Mice were trained in this protocol (Phase 2) for 6 to 10 days. During the weekend, water restriction was interrupted. Mice were weighed each morning before the first training session. Water droplet size was adjusted (ranging from 2 to 5*µ*L) depending on the number of rewards obtained. Care was taken to ensure the animal’s growth curve was maintained. Electrophysiological recordings were conducted either in task-naive animals that had not undergone the task or in mice that had reached a reliable level of performance, typically after 6 to 10 days of training (12 to 20 sessions). Following their last training session, water was reintroduced in the cages and the brain slice preparation was performed the following day.

### Behavioral Analysis

The quantification of the foraging behavior was done using the centroid positions and time derived from video frames recorded at 25 Hz and information recorded each time the mice transitioned from one trapezoid to the next one around a given tower (Schaffhauser et al., 2025). The trajectory of the mice was smoothed with a gaussian filter (σ = 1). The detection of the run from one spout turns was done in two steps (referred to as QTs). First, all the continuous bouts during which the mice ran without pause were detected. Pauses were defined as epochs during which the mouse’s speed was below 7 cm/s for at least 100 ms. Run bouts also had a minimal duration (at least 300 ms above 7 cm/s). Once all run epochs were detected, the initial and terminal positions of the mice were used to separate run around towers from other runs. Quarter-turns (QTs) around a given tower were defined as runs that were initiated and terminated in two adjacent trapezoids of a given tower. Basic kinematic parameters (duration, distance traveled, mean and peak speed, direction (CW or CCW), and reward status (rewarded or not)) were extracted for all QTs (Figures 2, 3, 4, S1, S2). Instantaneous speed was computed as the distance travelled between two frames divided by the time difference between the two frames. The obtained speed profiles were smoothed with a gaussian filter (σ = 1).

To evaluate the similarity between QTs trajectories, we computed the Fréchet distance, which is the minimum cord-length needed to connect a point moving along one trajectory to a point moving along another trajectory. Fréchet distance was computed across all possible pairs of QTs performed during a given session, separately for CW and CCW QTs (Figures 3, 4 and S2). This involved translating and rotating trajectories relative to a single point (a single corner of a single tower). First, the coordinates of the positions forming each QT (initially expressed in the arena reference frame) are transformed to be expressed relative to the specific tower corner around which the QT occurs. This step ensures that all QTs share a common reference point. Then, QT trajectories around the SW, NW and NE corners are rotated CCW by 90, 180 and 270 degrees, respectively. Thus, all the CW and CCW QTs performed during a session around different corners of different towers became aligned, allowing the quantification of Fréchet distances between pairs of CCW QTs and pairs of CW QTs. We use the median of all the Fréchet distances computed from all the pairs of CW and CCW QT as measure of the variability in their trajectory.

To quantify the extent of the mice’ engagement in reward collection, we computed their time spent within different zones of the arena (near the peripheral walls vs in the trapezes surrounding the towers).To quantify how much mice exploit a given tower, we then defined visits as sequences of QTs performed around a tower (Figures S1). A QT is considered as the start of a visit if the previous run epoch was not a QT in the same tower. For each visit, we computed the number of QTs, the number of rewarded QTs, the maximum number of rewards available and the starting time of the visit.

### Whisker Trimming

Whisker trimming was performed on 6 mice once task performance had stabilized. Mice completed a morning session (session 12 for 3 mice or session 14 for the other 3), which was followed by bilateral whisker trimming with scissors under isoflurane anesthesia (∼5 minutes of exposure). The sham group underwent identical isoflurane exposure without trimming. Five hours after sensory deprivation, an afternoon session was conducted. Behavioral parameters were compared before and after trimming to assess changes induced by sensory deprivation.

### Brain slices preparation and electrophysiology

All patch clamp recordings being performed in slices taken from the left hemisphere, their position, contralateral or ipsilateral to the whiskers stimulated by the towers during the task, changed according to the rewarded QTs. Thus, recordings were in the hemisphere contralateral to the stimulated whiskers when rewarded QTs had been performed in the counterclockwise direction (from the viewpoint of the camera that recorded the animals from below, i.e., clockwise direction from the top). Conversely, recordings were in the hemisphere ipsilateral to the stimulated whiskers (and contralateral to non-stimulated whiskers) when rewarded QTs had been clockwise (from below; counterclockwise from the top).

One day after their last session, mice were deeply anesthetized with isoflurane (4%) before cervical dislocation and decapitation. Parasagittal corticostriatal slices were prepared as previously described (Amroune et al., 2025). Slices used for LSPS contained barrels in the L4 of cortex, the globus pallidus, its external segment (GPe), the internal capsule, the ventral posteromedial nucleus of thalamus and the anterior hippocampus. Approximately 8 barrels (5-13) were visible in the slice. SPNs 50–150 *µ*m deep in the slice were visualized under infrared and fluorescent lights in a BX61WI microscope (Olympus) and patched with borosilicate electrodes (3–6 MΩ) and recorded in the voltage-clamp whole-cell configuration using a Multiclamp 700A amplifier (Axon Instrument, Molecular Devices). The holding membrane potential was – 80 mV. The intracellular solution contained (in mM) 128 Cs-methylsulfate, 4 MgCl2, 10 HEPES, 1 EGTA, 4 Na2ATP, 0.4 Na2GTP, 10 Na-phosphocreatine, 3 ascorbic acid; pH 7.25; 290-300 mOsm. Pyramidal neurons of the barrel cortex were recorded in the current-clamp mode, with an intracellular solution in which Cs-methylsulfate was replaced by K-methylsulfate. All experiments were performed at room temperature (21°C).

### LSPS with glutamate uncaging

LSPS was performed as described previously (Bureau et al., 2006). Recirculating (2 mL/min)ACSF solution contained (in mM): 0.2 MNI-caged glutamate (Tocris), 0.005 CPP [()-3-(2-carboxypiperazin-4-yl) propyl-1-phosphonic acid], 4 CaCl2, and 4 MgCl2. Focal photolysis of caged glutamate was achieved using a 2 ms pulse from a pulsed UV (355 nm) laser (DPSS Lasers Inc.) through a 0.16 NA 4× objective (Olympus). A 20 mW pulse was used for stimulating cortical neurons in the contraversive hemisphere, while a 25 mW pulse was employed for naive animals and for stimulation in the ipsiversive hemisphere to maintain excitation of cortical neurons at comparable levels. Ephus software for instrument control and acquisition was used (Suter et al., 2010). The stimulus pattern for mapping the corticostriatal projections was 464 positions spaced by 75 *µ*m on a 29 × 16 grid (2.1 × 1.1 mm) over barrel cortex. The corticostriatal slice and the LSPS grid were oriented in such a way that layer 5a was laid out horizontally. UV stimuli were applied every 700 ms and their successive positions on the LSPS grid were such as to maximize the time between stimulations of neighboring sites. Electrophysiological traces consisted of 100 ms baseline, a 450 ms window followed by a −5 mV / 100 ms test pulse. A minimum of 2 and up to 4 stimulations were performed at each site at several minutes intervals. Excitation profiles of pyramidal neurons were generated under similar conditions except that cells were recorded in current-clamp mode and glutamate was uncaged on a smaller 8 × 8 grid covering their soma and dendrites (50 *µ*m spacing; 350 × 350 *µ*m).

### Analysis of LSPS data

Synaptic input maps of neurons were constructed by taking the peak amplitude of EPSCs detected in a 50 ms time window starting at stimulation onset for each position in the LSPS grid. Measures were averaged across repetitions of stimulations (2-4). The threshold for EPSC detection was 3 standard deviations from baseline or 9.2 ± 0.4 pA. To disambiguate evoked responses from spontaneous activity, synaptic responses occurring less than 2 times across repetitions of maps were set to zero. To detect clusters of inputs in the LSPS map, we used the binary version of the map reporting the location of connected and non-connected sites (i.e., yielding EPSCs or none in the recorded SPN) and stacked it along its vertical axis. A cluster of inputs comprised 1 or more consecutive connected sites in the stacked map that was framed by 1 or more non-connected sites. Thus, a cluster here may include connected sites that were not adjacent on the vertical axis in the original map, and may combine synaptic inputs from different layers. The number of cortical columns was estimated based on the number of consecutive connected sites (ConsSites) in the stacked map and the mean width of a barrel in the slice, 150 *µ*m, which is equal to 2 connected sites in the map. Hence, the number of cortical columns = Σ (ceiling(ConsSites/2)).

Traces from current clamp recordings were analyzed in a 50 ms time window at stimulus onset to count the number of action potentials (APs) elicited upon glutamate uncaging at every site and in the entire stimulation grid.

### Statistical Analysis

Median and 25-75th percentiles are shown in the Figures. N and n are the numbers of animals and neurons, respectively. Statistical analyses were performed with Python scripts using permutation tests (n = 10000 iterations), and Wilcoxon signed ranks test for paired comparison. Results were considered statistically significant for a p-value < 0.05, except for multiple comparisons where we divided 0.05 by the number of comparisons. Differences in layer input proportions between the groups were assessed using a permutation-based approach. Each cell was treated as an independent observation and represented by a binary vector indicating the presence or absence of input across layers. For each layer, the observed difference in mean input proportion between groups was computed. To generate the null distribution, group labels were randomly permuted across cells. For each permutation (n = 1,000), mean layer proportions were recalculated for the two permuted groups and the difference between them was recorded. The observed group difference was then compared to the permutation distribution. A permutation-based confidence interval was defined using the 2.5th and 97.5th percentiles of all the permuted differences. Differences were considered significant (taking in account multiple comparisons) when the observed values fell outside this confidence interval.

## Acknowledgements

This work was supported by funding from the Institut National de la Santé et de la Recherche Médicale and a grant from the Agence Nationale de la Recherche (Corticostriatal, ANR-20-CE16-0002). K. Amroune was supported by fellowships from the Ministère de l’Enseignement Supérieur et de la Recherche and from the French government under the “France 2030” program via A*Midex (Initiative d’Excellence d’Aix-Marseille Université, AMX-19-IET-004) and ANR funding (ANR-17-EURE-0029) and a grant from Aix-Marseille University (AMX-22-RE-AB-007/Neuradventure). T. Morvan was supported by the Fondation pour la Recherche Médicale (FDT202204014828). We thank Mathias Lechelon and Robert Martinez for help with the behavioral set-up, Tom Orgelet-Lacomme for the python codes to analyse the behavior of mice and the interns who collaborated on this project (Cloe Alcaraz, Ella Flores, Alexy Louis, Lola Michaud, and Océane Pierrot) as well as the staff of the animal and genotyping facilities at INMED. We are grateful to the members of the CBGB group for their assistance and support.

## Authors contributions

Kenza Amroune: Conceptualization; Methodology; Software; Formal analysis; Investigation; Visualization; Writing – original draft; Writing – review & editing.

Maud Schaffhauser: Methodology; Writing – review & editing.

Thomas Morvan: Methodology; Software; Formal analysis; Writing – review & editing. Ingrid Bureau: Conceptualization; Methodology; Software; Formal analysis; Visualization, Data curation; Writing – review & editing; Supervision; Project administration; Funding acquisition.

David Robbe: Conceptualization; Methodology; Software; Formal analysis; Visualization, Data curation; Writing – original draft; Writing – review & editing; Supervision; Project administration; Funding acquisition.

## Notes

### Competing Interest Statement

The authors have declared no competing interest.

## References

Ade KK, Wan Y, Chen M, Gloss B, Calakos N (2011) An Improved BAC Transgenic Fluorescent Reporter Line for Sensitive and Specific Identification of Striatonigral Medium Spiny Neurons. Front Syst Neurosci 5:32.

Amroune K, Fontolan L, Baude A, Robbe D, Bureau I (2025) Sparse innervation and local heterogeneity in the vibrissal corticostriatal projection. eLife 14:RP106621.

Barbera G, Liang B, Zhang L, Gerfen CR, Culurciello E, Chen R, Li Y, Lin D-T (2016) Spatially Compact Neural Clusters in the Dorsal Striatum Encode Locomotion Relevant Information. Neuron 92:202–213.

Barnes TD, Kubota Y, Hu D, Jin DZ, Graybiel AM (2005) Activity of striatal neurons reflects dynamic encoding and recoding of procedural memories. Nature 437:1158–1161.

Bay Kønig A, Ciriachi C, Gether U, Rickhag M (2019) Chemogenetic Targeting of Dorsomedial Direct-pathway Striatal Projection Neurons Selectively Elicits Rotational Behavior in Mice. Neuroscience 401:106–116.

Bertran-Gonzalez J, Bosch C, Maroteaux M, Matamales M, Hervé D, Valjent E, Girault J-A (2008) Opposing patterns of signaling activation in dopamine D1 and D2 receptor-expressing striatal neurons in response to cocaine and haloperidol. J Neurosci Off J Soc Neurosci 28:5671–5685.

Bonnavion P, Varin C, Fakhfouri G, Martinez Olondo P, De Groote A, Cornil A, Lorenzo Lopez R, Pozuelo Fernandez E, Isingrini E, Rainer Q, Xu K, Tzavara E, Vigneault E, Dumas S, De Kerchove d’Exaerde A, Giros B (2024) Striatal projection neurons coexpressing dopamine D1 and D2 receptors modulate the motor function of D1- and D2-SPNs. Nat Neurosci 27:1783–1793.

Bureau I, von Saint Paul F, Svoboda K (2006) Interdigitated paralemniscal and lemniscal pathways in the mouse barrel cortex. PLoS Biol 4:e382.

Cao J, Dorris DM, Meitzen J (2018) Electrophysiological properties of medium spiny neurons in the nucleus accumbens core of prepubertal male and female Drd1a-tdTomato line 6 BAC transgenic mice. J Neurophysiol 120:1712–1727.

Cregg JM, Sidhu SK, Leiras R, Kiehn O (2024) Basal ganglia–spinal cord pathway that commands locomotor gait asymmetries in mice. Nat Neurosci 27:716–727.

Cruz BF, Guiomar G, Soares S, Motiwala A, Machens CK, Paton JJ (2022) Action suppression reveals opponent parallel control via striatal circuits. Nature 607:521–526.

Cui G, Jun SB, Jin X, Pham MD, Vogel SS, Lovinger DM, Costa RM (2013) Concurrent activation of striatal direct and indirect pathways during action initiation. Nature 494:238–242.

Dhawale AK, Miyamoto YR, Smith MA, Ölveczky BP (2019) Adaptive Regulation of Motor Variability. Curr Biol 29:3551–3562.e7.

Dhawale AK, Wolff SBE, Ko R, Ölveczky BP (2021) The basal ganglia control the detailed kinematics of learned motor skills. Nat Neurosci 24:1256–1269.

Dunovan K, Verstynen T (2016) Believer-Skeptic Meets Actor-Critic: Rethinking the Role of Basal Ganglia Pathways during Decision-Making and Reinforcement Learning. Front Neurosci 10.

Enoksson T, Bertran-Gonzalez J, Christie MJ (2012) Nucleus accumbens D2- and D1-receptor expressing medium spiny neurons are selectively activated by morphine withdrawal and acute morphine, respectively. Neuropharmacology 62:2463–2471.

Hidalgo-Balbuena AE, Luma AY, Pimentel-Farfan AK, Peña-Rangel T, Rueda-Orozco PE (2019) Sensory representations in the striatum provide a temporal reference for learning and executing motor habits. Nat Commun 10:4074.

Hikida T, Kimura K, Wada N, Funabiki K, Nakanishi S (2010) Distinct Roles of Synaptic Transmission in Direct and Indirect Striatal Pathways to Reward and Aversive Behavior. Neuron 66:896–907.

Humphries MD, Khamassi M, Gurney K (2012) Dopaminergic Control of the Exploration-Exploitation Trade-Off via the Basal Ganglia. Front Neurosci 6:9.

Hwang F-J, Roth RH, Wu Y-W, Sun Y, Kwon DK, Liu Y, Ding JB (2022) Motor learning selectively strengthens cortical and striatal synapses of motor engram neurons. Neuron 0.

Jacob V, Petreanu L, Wright N, Svoboda K, Fox K (2012) Regular Spiking and Intrinsic Bursting Pyramidal Cells Show Orthogonal Forms of Experience-Dependent Plasticity in Layer V of Barrel Cortex. Neuron 73:391–404.

Jin X, Costa RM (2010) Start/stop signals emerge in nigrostriatal circuits during sequence learning. Nature 466:457–462.

Jurado-Parras M-T, Safaie M, Sarno S, Louis J, Karoutchi C, Berret B, Robbe D (2020) The Dorsal Striatum Energizes Motor Routines. Curr Biol CB 30:4362–4372.e6.

Kim N, Barter JW, Sukharnikova T, Yin HH (2014) Striatal firing rate reflects head movement velocity. Eur J Neurosci 40:3481–3490.

Klaus A, Martins GJ, Paixao VB, Zhou P, Paninski L, Costa RM (2017) The Spatiotemporal Organization of the Striatum Encodes Action Space. Neuron 95:1171–1180.e7.

Klug JR, Yan X, Hoffman H, Engelhardt MD, Osakada F, Callaway EM, Jin X (2025) Asymmetric cortical projections to striatal direct and indirect pathways distinctly control actions Ding J, Wassum KM, eds. eLife 12:RP92992.

Koralek AC, Costa RM, Carmena JM (2013) Temporally Precise Cell-Specific Coherence Develops in Corticostriatal Networks during Learning. Neuron 79:865–872.

Koralek AC, Jin X, Long II JD, Costa RM, Carmena JM (2012) Corticostriatal plasticity is necessary for learning intentional neuroprosthetic skills. Nature 483:331–335.

Kravitz AV, Freeze BS, Parker PRL, Kay K, Thwin MT, Deisseroth K, Kreitzer AC (2010) Regulation of parkinsonian motor behaviours by optogenetic control of basal ganglia circuitry. Nature 466:622–626.

Kravitz AV, Tye LD, Kreitzer AC (2012) Distinct roles for direct and indirect pathway striatal neurons in reinforcement. Nat Neurosci 15:816–818.

Kreitzer AC, Malenka RC (2008) Striatal Plasticity and Basal Ganglia Circuit Function. Neuron 60:543–554.

Lemke SM, Ramanathan DS, Darevksy D, Egert D, Berke JD, Ganguly K (2021) Coupling between motor cortex and striatum increases during sleep over long-term skill learning. eLife 10:e64303.

Lemke SM, Ramanathan DS, Guo L, Won SJ, Ganguly K (2019) Emergent modular neural control drives coordinated motor actions. Nat Neurosci 22:1122–1131.

Lowet AS, Zheng Q, Meng M, Matias S, Drugowitsch J, Uchida N (2025) An opponent striatal circuit for distributional reinforcement learning. Nature 639:717–726.

Markowitz JE, Gillis WF, Beron CC, Neufeld SQ, Robertson K, Bhagat ND, Peterson RE, Peterson E, Hyun M, Linderman SW, Sabatini BL, Datta SR (2018) The Striatum Organizes 3D Behavior via Moment-to-Moment Action Selection. Cell 174:44–58.e17.

Mink JW (1996) The basal ganglia: focused selection and inhibition of competing motor programs. Prog Neurobiol 50:381–425.

Mizes KGC, Lindsey J, Escola GS, Ölveczky BP (2023) Dissociating the contributions of sensorimotor striatum to automatic and visually guided motor sequences. Nat Neurosci 26:1791–1804.

Nonomura S, Nishizawa K, Sakai Y, Kawaguchi Y, Kato S, Uchigashima M, Watanabe M, Yamanaka K, Enomoto K, Chiken S, Sano H, Soma S, Yoshida J, Samejima K, Ogawa M, Kobayashi K, Nambu A, Isomura Y, Kimura M (2018) Monitoring and Updating of Action Selection for Goal-Directed Behavior through the Striatal Direct and Indirect Pathways. Neuron 99:1302–1314.e5.

O’Connor DH, Clack NG, Huber D, Komiyama T, Myers EW, Svoboda K (2010) Vibrissa-based object localization in head-fixed mice. J Neurosci Off J Soc Neurosci 30:1947–1967.

Panigrahi B, Martin KA, Li Y, Graves AR, Vollmer A, Olson L, Mensh BD, Karpova AY, Dudman JT (2015) Dopamine Is Required for the Neural Representation and Control of Movement Vigor. Cell 162:1418–1430.

Park J, Polidoro P, Fortunato C, Arnold J, Mensh B, Gallego JA, Dudman JT (2025) Conjoint specification of action by neocortex and striatum. Neuron 113:620–636.e6.

Parker JG, Marshall JD, Ahanonu B, Wu Y-W, Kim TH, Grewe BF, Zhang Y, Li JZ, Ding JB, Ehlers MD, Schnitzer MJ (2018) Diametric neural ensemble dynamics in parkinsonian and dyskinetic states. Nature 557:177–182.

Peters, Fabre JMJ, Steinmetz NA, Harris KD, Carandini M (2021) Striatal activity topographically reflects cortical activity. Nature 591:420–425.

Reiner A, Jiao Y, Del Mar N, Laverghetta AV, Lei WL (2003) Differential morphology of pyramidal tract-type and intratelencephalically projecting-type corticostriatal neurons and their intrastriatal terminals in rats. J Comp Neurol 457:420–440.

Rosselet C, Fieschi M, Hugues S, Bureau I (2011) Associative learning changes the organization of functional excitatory circuits targeting the supragranular layers of mouse barrel cortex. Front Neural Circuits 4:126.

Rueda-Orozco PE, Robbe D (2015) The striatum multiplexes contextual and kinematic information to constrain motor habits execution. Nat Neurosci 18:453–460.

Sachidhanandam S, Sreenivasan V, Kyriakatos A, Kremer Y, Petersen CCH (2013) Membrane potential correlates of sensory perception in mouse barrel cortex. Nat Neurosci 16:1671–1677.

Safaie M, Jurado-Parras M-T, Sarno S, Louis J, Karoutchi C, Petit LF, Pasquet MO, Eloy C, Robbe D (2020) Turning the body into a clock: Accurate timing is facilitated by simple stereotyped interactions with the environment. Proc Natl Acad Sci 117:13084–13093.

Sales-Carbonell C, Taouali W, Khalki L, Pasquet MO, Petit LF, Moreau T, Rueda-Orozco PE, Robbe D (2018) No Discrete Start/Stop Signals in the Dorsal Striatum of Mice Performing a Learned Action. Curr Biol 28:3044–3055.e5.

Santos FJ, Oliveira RF, Jin X, Costa RM (2015) Corticostriatal dynamics encode the refinement of specific behavioral variability during skill learning Kiehn O, ed. eLife 4:e09423.

Schaffhauser M, Orjollet-Lacomme T, Amroune K, Morvan T, Fortoul A, Lechelon M, Robbe D (2025) A Novel Foraging Task Reveals Cognitive and Motor Processes Underlying Behavioral Flexibility. bioRxiv:2025.09.09.675194.

Shepherd GMG (2013) Corticostriatal connectivity and its role in disease. Nat Rev Neurosci 14:278–291.

Shepherd GMG, Pologruto TA, Svoboda K (2003) Circuit analysis of experience-dependent plasticity in the developing rat barrel cortex. Neuron 38:277–289.

Sippy T, Chaimowitz C, Crochet S, Petersen CCH (2021) Cell Type-Specific Membrane Potential Changes in Dorsolateral Striatum Accompanying Reward-Based Sensorimotor Learning. Function 2:zqab049.

Sippy T, Lapray D, Crochet S, Petersen CCH (2015) Cell-Type-Specific Sensorimotor Processing in Striatal Projection Neurons during Goal-Directed Behavior. Neuron 88:298–305.

Suter BA, O’Connor T, Iyer V, Petreanu LT, Hooks BM, Kiritani T, Svoboda K, Shepherd GMG (2010) Ephus: multipurpose data acquisition software for neuroscience experiments. Front Neural Circuits 4:100.

Tai L-H, Lee AM, Benavidez N, Bonci A, Wilbrecht L (2012) Transient stimulation of distinct subpopulations of striatal neurons mimics changes in action value. Nat Neurosci 15:1281–1289.

Tecuapetla F, Matias S, Dugue GP, Mainen ZF, Costa RM (2014) Balanced activity in basal ganglia projection pathways is critical for contraversive movements. Nat Commun 5:4315.

Thibault D, Loustalot F, Fortin GM, Bourque M-J, Trudeau L-É (2013) Evaluation of D1 and D2 dopamine receptor segregation in the developing striatum using BAC transgenic mice. PloS One 8:e67219.

Turrigiano GG (2008) The Self-Tuning Neuron: Synaptic Scaling of Excitatory Synapses. Cell 135:422–435.

Wall NR, De La Parra M, Callaway EM, Kreitzer AC (2013) Differential innervation of direct-and indirect-pathway striatal projection neurons. Neuron 79:347–360.

Wen W, Turrigiano GG (2024) Keeping Your Brain in Balance: Homeostatic Regulation of Network Function. Annu Rev Neurosci 47:41–61.

Wise SP, Jones EG (1977) Cells of origin and terminal distribution of descending projections of the rat somatic sensory cortex. J Comp Neurol 175:129–157.

Xiong Q, Znamenskiy P, Zador AM (2015) Selective corticostriatal plasticity during acquisition of an auditory discrimination task. Nature 521:348–351.

Yamashita T, Petersen CC (2016) Target-specific membrane potential dynamics of neocortical projection neurons during goal-directed behavior. eLife 5:e15798.

Yamashita T, Vavladeli A, Pala A, Galan K, Crochet S, Petersen SSA, Petersen CCH (2018) Diverse Long-Range Axonal Projections of Excitatory Layer 2/3 Neurons in Mouse Barrel Cortex. Front Neuroanat 12:33.

Yin HH, Mulcare SP, Hilário MR, Clouse E, Holloway T, Davis MI, Hansson AC, Lovinger DM, Costa RM (2009) Dynamic reorganization of striatal circuits during the acquisition and consolidation of a skill. Nat Neurosci 12:333–341.

Yttri EA, Dudman JT (2016) Opponent and bidirectional control of movement velocity in the basal ganglia. Nature 533:402–406.

Zareian B, Lam A, Zagha E (2023) Dorsolateral Striatum is a Bottleneck for Responding to Task-Relevant Stimuli in a Learned Whisker Detection Task in Mice. J Neurosci Off J Soc Neurosci 43:2126–2139.

Zheng Q, Liu Y, Huang Y, Cao J, Wang X, Yu J (2025) The Role of Striatum in Controlling Waiting during Reactive and Self-Timed Behaviors. J Neurosci 45.

